# Anatomical and functional maturation of the mid-gestation human intestine

**DOI:** 10.1101/2022.08.02.501641

**Authors:** Lori B. Dershowitz, Li Li, Anca M. Pasça, Julia A. Kaltschmidt

## Abstract

Gastrointestinal (GI) maturation is a key determinant of survival for extremely preterm infants. The enteric nervous system (ENS) controls GI motility, and immature GI motility limits enteral feeding and causes severe health complications.^1^ Due to the significant challenges in obtaining and studying human fetal tissue, little is known about when the human ENS becomes mature enough to carry out vital functions. Here we define the progressive anatomical maturation of the human fetal ENS and analyze GI motility in the second trimester of *in utero* development. We identify substantial structural changes in the ENS including the emergence of striped neuronal cytoarchitecture and a shift in the representation of excitatory and inhibitory neurons. We further analyze and pharmacologically manipulate GI motility in freshly collected human fetal intestines, which, to our knowledge, is a first functional analysis of intact human fetal organs *ex vivo*. We find that the ENS influences GI motility beginning at 21 postconceptional weeks (PCW), the earliest reported evidence of neurogenic GI motility. Our study provides unprecedented insight into human fetal ENS development, foundational knowledge which facilitates comparisons with common animal models to advance translational disease investigations and testing of pharmacological agents to enhance GI motility in prematurity.

## Introduction

GI immaturity is a major source of morbidity and mortality amongst extremely preterm infants born at less than 28PCW. These patients are fed enterally despite immature GI motility and are therefore at high risk of feeding intolerance, functional intestinal obstruction, and subsequent overwhelming infection. Treatments to alleviate GI immaturity are lacking. Clinical trials on extremely preterm infants are highly challenging, and attempts to use adult prokinetic agents such as erythromycin have failed to treat feeding intolerance in the preterm population.^1,2^ Therefore, a fundamental understanding of mid-gestation GI physiology is critical for developing therapies.

The ENS, the autonomous branch of the peripheral nervous system, resides within the intestinal walls and controls GI function. Studies in mice have shown that the ENS gradually reorganizes during late embryonic and early postnatal ages,^3^ which corresponds to the emergence of GI motility.^4,5^ However, our understanding of human ENS development, structure, and function *in utero* remains extremely limited.

Whole-intestine single-cell RNA sequencing (scRNAseq) has revealed that enteric neurons are present in the intestine as early as 6PCW and that genes indicative of functionally distinct enteric neuron classes are expressed during the first trimester.^6–8^ However, these studies include limited second trimester data and provide minimal information about ENS structure. Similarly, GI motility assessments have been restricted to manometry studies in preterm and term infants from 28-42PCW. These recordings primarily identified random and non-propagating GI activity at preterm ages and did not detect migrating motor complexes, a form of neuronally mediated GI motility, until 37PCW.^9,10^

We provide the first description of human fetal ENS structural development and *ex vivo* GI motility during the second trimester, between 14-23PCW. This new knowledge of ENS development from a period that closely precedes extremely preterm birth is essential for understanding the pathophysiology of severe GI disorders and for guiding new enteral feeding regimens and pharmacologic interventions for GI immaturity.

## Results

In cross section, the intestinal wall contains five distinct layers: mucosa, submucosal plexus (SMP) of the ENS, circular muscle (CM), myenteric plexus (MP) of the ENS, and longitudinal muscle (LM). To identify morphological stages of ENS development in the second trimester of pregnancy, we collected fresh *ex vivo* human intestines from non-diseased fetuses between 14-23PCW (Fig.1a). For analyses, we subdivided the intestine into five anatomically and functionally distinct regions (duodenum, jejunum, ileum, proximal colon, and distal colon) (Fig. 1a). Immunolabeling of jejunum and proximal colon cross sections with pan-neuronal markers PGP9.5 and HuC/D revealed a uniform MP with few neurons in the SMP at 14PCW (Fig.1b, Supplementary Fig.1a). By 23PCW, enteric neurons have condensed into discrete ganglia in both the SMP and MP, and the CM and mucosa contain a greater density of neuronal fibers, which may arise from either extrinsic or intrinsic innervation (Fig.1b, Supplementary Fig.1a). We also observed substantial changes in mucosa morphology and increased CM and LM thickness from 14 to 23PCW (Fig.1b, Supplementary Fig.1a). Thus, human intestinal and ENS morphology develop substantially during mid-gestation.

**Figure 1:**
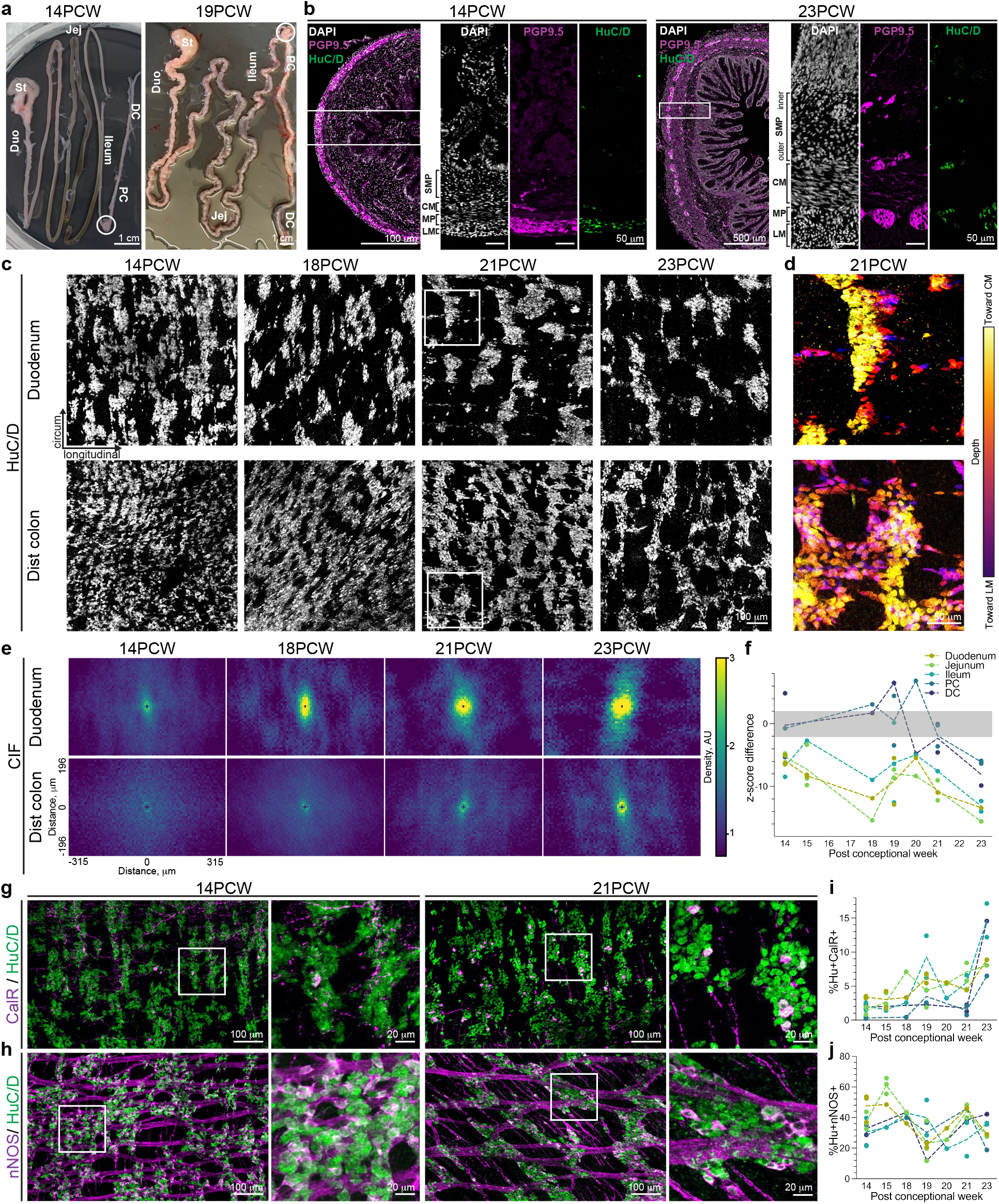
Anatomical development of the mid-gestation human intestine and ENS. (**a**) Images of 14 and 19PCW human fetal intestinal samples with regional subdivisions indicated. White circles denote the appendix. (**b**) Representative images of cross sections from the proximal colon at 14 and 23PCW with immunohistochemical (IHC) labeling against DAPI (white), pan-neuronal markers PGP9.5 (magenta) and HuC/D (green). White boxes indicate locations of higher magnitude insets. Brackets denote location of the submucosal plexus (SMP), circular muscle (CM), myenteric plexus (MP), and longitudinal muscle (LM). (**c**) Representative images of HuC/D labeling in MP wholemounts of the duodenum (top) and distal colon (bottom) over gestational time. Scale bar applies to all images. (**d**) Depth projection of HuC/D+ neurons in the 21PCW MP as indicated by white boxes in (c). Yellow: closer to CM; purple: closer to LM. Scale bar applies to all images. (**e**) Conditional intensity function (CIF) plots derived from HuC/D labeling in MP wholemounts of the duodenum (top) and distal colon (bottom) over gestational time. Yellow: high probability density; blue: low. Axes apply to all panels. (**f**) z-score difference between data and mean of synthetically generated pseudorandom values in all intestinal regions over gestational time. Gray bar indicates random value range. n = 1-3. (**g**,**h**) HuC/D (green) and excitatory marker CalR (**g**, purple) or inhibitory marker nNOS (**h**, purple) in the jejunum MP at 14 and 21PCW. White boxes indicate locations of higher magnitude insets. (**i**,**j**) Proportion of total HuC/D+ neurons positive for CalR (**i**) or nNOS (**j**) (mean ± SEM) over gestational time. n = 1-3. Scale bars as indicated. AU, arbitrary units; Duo, duodenum; DC, distal colon; Jej, jejunum; PC, proximal colon; PCW, postconceptional week.

In mouse, the embryonic and neonatal MP undergoes progressive reorganization from a random array into circumferentially oriented neuronal stripes, which define the locations of enteric ganglia.^3^ To determine whether the human fetal ENS undergoes similar structural reorganization, we visualized enteric neurons in MP wholemounts at ages spanning 14-23PCW (Fig.1c). In the duodenum MP, enteric neurons reside in circumferentially oriented neuronal stripes as early as 14PCW (Fig.1c). In the distal colon MP, distinct neuronal stripes are not evident until 21PCW (Fig.1c). The emergence of neuronal stripes is not due to programmed cell death as labeling with apoptotic marker Caspase3 identified few apoptotic neurons (Supplementary Fig.1b). To validate these observations with a computational non-biased approach, we generated spatial probability maps of neuronal location within our tissue (Fig.1e). Z-score comparisons between neuronal data and synthetically generated pseudorandom data further demonstrated that neuronal organization becomes non-random and more clustered in the small intestine prior to the colon (Fig.1c,f).^3^ Furthermore, this analysis identified neurons within either of two distinct positions: (1) neurons within stripes, which reside close to the CM and (2) bridging neurons, which reside closer to the LM (Fig.1c-e). Together, they form a grid-like structure. We primarily focused on MP development due to its role in GI motility, but we also noted structural changes in SMP wholemounts including increased ganglia size from 15 to 23PCW (Supplementary Fig.1c). Taken together, these results demonstrate that the mid-gestation fetal MP undergoes substantial reorganization, which occurs in the small intestine prior to the colon.

The basic GI motility circuit in rodents contains four functionally distinct neuron classes.^11^ We assessed the quantity and distribution of these classes across human MP development. We labeled MP wholemounts with antibodies against calretinin (CalR), neuronal nitric oxide synthase (nNOS), neurofilament-M (NF-M), and somatostatin (SST), which broadly define excitatory motor neurons, inhibitory motor neurons, sensory neurons, and interneurons, respectively (Fig.1g-j, Supplementary Fig.1d-g).^11–13^ All markers were detected as early as 14PCW except SST, which was not detected until 15PCW (Fig.1g-j, Supplementary Fig.1d-g). The percentage of CalR+HuC/D+ neurons increased over gestational time in all intestinal regions while the percentage of nNOS+HuC/D+ neurons appeared to decrease over time, although individual regions did not all share this overarching trend (Fig.1g-j). This result is consistent with reported transient nNOS expression in a subset of mouse enteric neurons embryonically.^14^ Labeling with NF-M did not reveal an obvious trend across gestational time or region, and SST+HuC/D+ neurons remained scarce until 21PCW (Supplementary Fig.1d-g). Thus, the mid-gestation human MP contains distinct neuron classes with apparent increasing representation of excitatory subtypes and decreasing representation of inhibitory subtypes.

Finally, we assessed how these organizational changes relate to human GI motility development. We established a GI motility monitor used in guinea pig and mouse^15–17^ for *ex vivo* motility analysis of the second trimester human small intestine (Supplementary movies 1-8). Using spatiotemporal maps (STMs), which capture changes in intestinal diameter over time, we identified contractions that travel both proximally and distally from initiation points at 14 and 15PCW (Fig.2a,b, Supplementary movies 1,2) and increase in frequency until 20PCW (Fig.2d). At 20PCW, we occasionally observed bias toward distal progression of contractions (Fig.2a, Supplementary movie 3). Notably, at 21PCW we ceased to observe bidirectional events and instead found evidence of contractions that propagate exclusively from proximal to distal (Fig.2a,b, Supplementary movie 4). To determine whether these GI motility patterns are primarily myogenic ripples^5^ or neuronally mediated contractions, we inhibited neuronal activity using tetrodotoxin (TTX), a non-specific sodium channel blocker. At 14 and 15PCW, TTX treatment affected contraction speed and frequency in some samples but did not alter the overall shape of GI motility patterns (Fig.2c-e, Supplementary movies 5,6). At 21PCW, TTX substantially altered GI motility and disrupted the exclusively distal direction of contractions (Fig.2c, Supplementary movies 7,8). Collectively, these results suggest that GI motility is limited to myogenic ripples at 14-15PCW and that the ENS may first influence GI motility at 21PCW.

**Figure 2:**
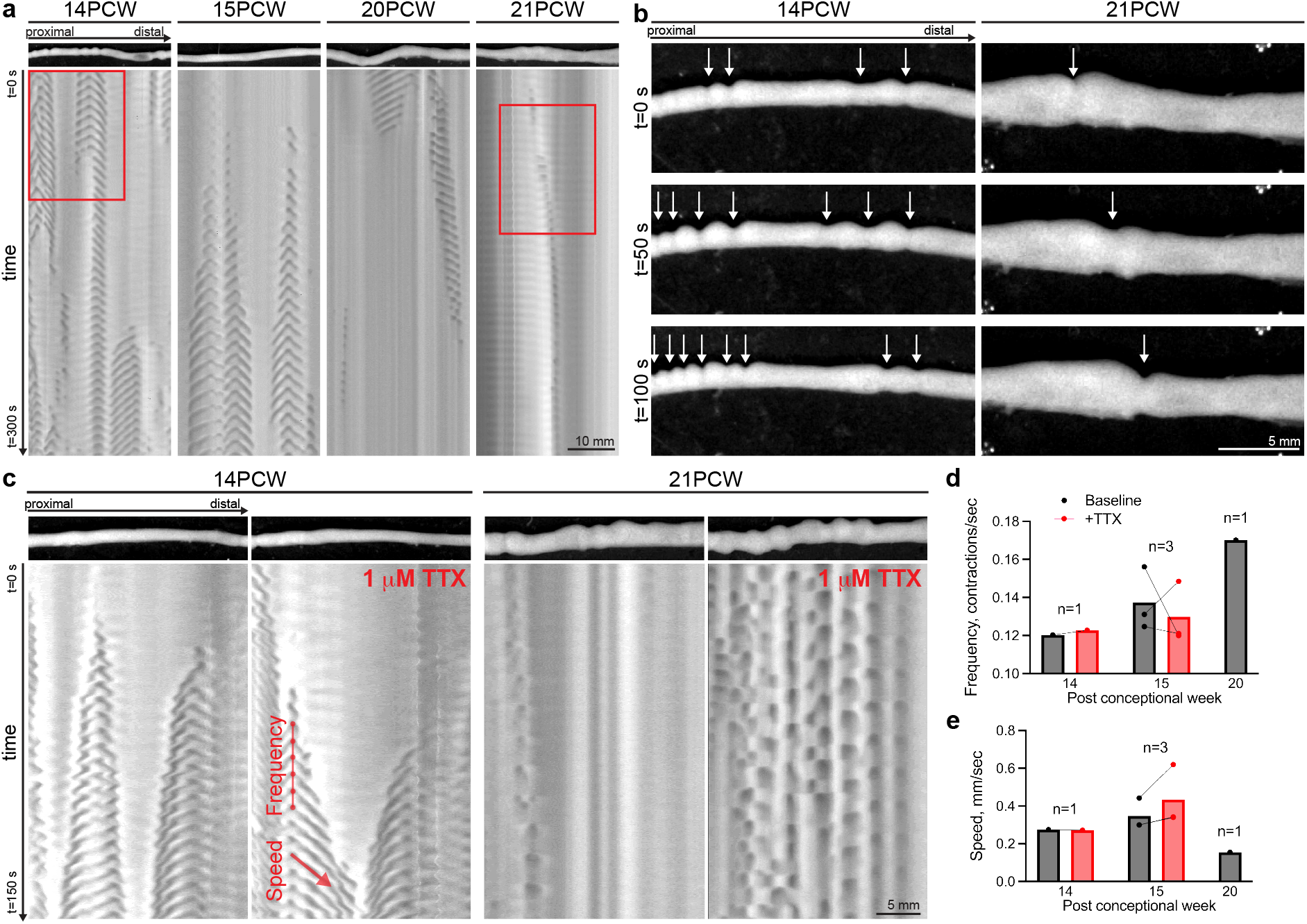
GI motility in the 2^nd^ trimester human small intestine assessed *ex vivo*. (**a**) Spatiotemporal maps (STMs) with accompanying images of GI motility in the human fetal jejunum at 14, 15, 20, and 21PCW. Dark gray: decreased intestinal diameter; light gray: increased. (**b**) Images from video recordings over three timepoints indicated by red boxes in (a) from 14 and 21PCW. White arrows indicate contractions. Scale bar applies to all images. (**c**) STMs with representative images of GI motility in the human fetal duodenum at 14 and 21PCW at baseline (left panels) and with the addition of 1 μM tetrodotoxin (TTX, right panels). Dark gray: decreased intestinal diameter; light gray: increased. Scale bar applies to all images. (**d**) Frequency of contractions as indicated in (c) at baseline (black bars, 14, 15, 20 PCW) and with TTX (red bars, 14 and 15PCW). n = 1-3. (**e**) Speed of contractions as indicated in (c) at baseline (black bars, 14, 15, 20 PCW) and with TTX (red bars, 14 and 15PCW). n = 1-3.

## Discussion

Here, we provide the first characterization of the anatomical and functional development of the human fetal ENS with a focus on second trimester *in utero* development. First, we identified emerging striped neuronal cytoarchitecture of the human MP in the duodenum at 14PCW and in the distal colon at 21PCW (Fig.1c-f), a result similar to mouse MP development at late embryonic and early neonatal ages.^3^ Interestingly, we also observed a neuron grid (Fig.1c-e), a more complex macrostructure than observed in mouse.^3^ Understanding this structural progression is of major clinical significance for neonatology and maternal-fetal medicine because this process may be disrupted in cases of utero-placental insufficiency, which results in de-prioritization of fetal intestinal development.^18^

Second, our findings suggest that the representation of CalR+ excitatory neurons increases relative to nNOS+ inhibitory neurons in the human MP during the second trimester (Fig.1g-j). This result is an important clinical benchmark because decreased excitatory/inhibitory neuron ratios have been observed in the ganglionated portion of the intestine of mouse models and individuals with Hirschsprung’s disease.^19^

Third, we establish the first *ex vivo* assay of function in an intact human fetal organ. Our *ex vivo* GI motility analysis identified myogenic ripples at 14PCW that increase in frequency over gestational time and found evidence of neuronally mediated GI motility at 21PCW (Fig.2a-e; Supplementary movies 1-8). While future studies will need to include more gestational ages, our findings suggest incipient maturation of the ENS motility circuit beginning at 21PCW, an age potentially amenable to clinical interventions to enhance ENS function in preterm infants. Collectively, our studies provide unprecedented insights into the mid-gestation human ENS that allow for direct comparisons between human and mouse ENS development and may improve use of animal models for studying GI diseases with an *in utero* onset. Further, we demonstrate that the *ex vivo* platform can be used with pharmacologic agents thus enabling noninvasive screening of drugs that enhance GI motility with the long-term goal of reducing feeding intolerance and serious complications including intestinal perforation and sepsis.

## Methods

### Human tissue collection

De-identified intestinal tissue samples were obtained after elective pregnancy termination at Stanford University School of Medicine under an approved protocol through the Research Compliance Office at Stanford University. Tissue postconceptional week (PCW) was estimated based on ultrasound appearance and last menstrual period prior to pregnancy termination. For histology experiments, all intestinal tissue was immediately placed in ice-cold PBS. For functional experiments, tissue was immediately placed in carbogenated (95% CO_2_, 5% O_2_) Krebs solution (pH 7.4 containing (in mmol/l): 117 NaCl, 4.7 KCl, 3.3 CaCl_2_ (2H_2_O), 1.5 MgCl_2_ (6H_2_O), 25 NaHCO_3_, 1.2 NaH_2_PO_4_ and 11 Glucose) at 37°C.

### Immunohistochemistry

For histological assessment of intestinal and enteric nervous system (ENS) structure, intestinal tissue was identified and dissected from surrounding organs, and the mesentery cut away. Based on tissue coloration and position relative to anatomical landmarks including the stomach and appendix (Fig.1a), the intestines were subdivided into stomach, duodenum, jejunum, ileum, proximal colon, and distal colon. For each region, a representative piece of tissue was collected. The length of each tissue piece varied by age, from ∼4 cm in length at 14PCW and ∼10 cm in length at 23PCW.

For cross sections, tissue was fixed as a tube with each end pinned on Sylgard 170 in 4% PFA for 2 hours at 4 °C while shaking. Tissue was placed in 30% sucrose overnight, embedded in OCT, and stored at -80 °C. Tissue sections were cut at 12mm thickness with a Leica cryostat CM3050 S. After being allowed to dry for 30 minutes, slides were rehydrated with 3 washes in ice-cold PBS and placed in blocking solution with 0.5% donkey serum for 1 hour at room temperature. Slides were processed with a Streptavidin/Biotin Blocking Kit (Vector Laboratories) per package instructions and incubated in primary antibodies (see Table 1) diluted in PBT (PBS, 1% BSA, 0.1% Triton X-100) overnight at 4 °C. Slides were washed 3 times in ice-cold PBS, incubated in PBT containing secondary antibodies (see Table 2) for 2 hours at room temperature, and washed 3 times in ice-cold PBS. Slides were rinsed in ddH_2_O and coverslipped with Fluoromount-G (Southern Biotech).

**Table 1:** Primary antibodies for immunohistochemistry

| Antibody | Host | Dilution | Source | Catalogue No. | RRID |
| --- | --- | --- | --- | --- | --- |
| Calretinin | Goat | 1:8000 | Swant | CG1 | AB_10000342 |
| Caspase3 | Rabbit | 1:500 | Abcam | AB2302 | AB_302962 |
| HuC/D | Mouse, biotinylated | 1:500 | ThermoFisher | A-21272 | AB_2535822 |
| Ki67 | Rabbit | 1:1000 | Abcam | ab15580 | AB_443209 |
| Neurofilament-M | Rabbit | 1:2000 | Millipore | AB1987 | AB_91201 |
| nNOS | Rabbit | 1:2000 | Sigma-Aldrich | N7280 | AB_260796 |
| PGP9.5 | Guinea pig | 1:1000 | Abcam | ab10410 | AB_287150 |
| Somatostatin | Rat | 1:500 | Millipore | MAB354 | AB_2255365 |

**Table 2:**
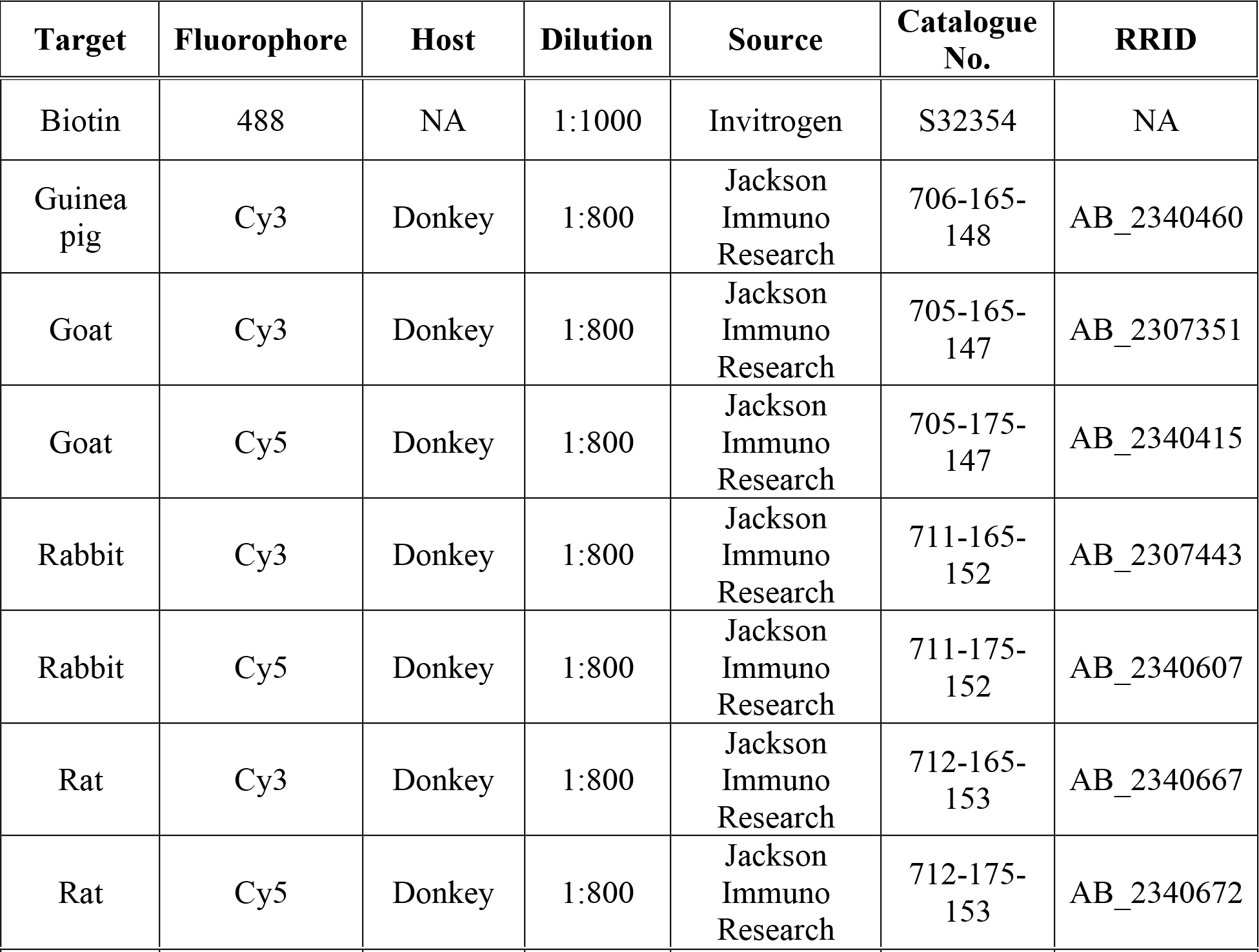
Secondary antibodies for immunohistochemistry

| Target | Fluorophore | Host | Dilution | Source | Catalogue No. | RRID |
| --- | --- | --- | --- | --- | --- | --- |
| Biotin | 488 | NA | 1:1000 | Invitrogen | S32354 | NA |
| Guinea pig | Cy3 | Donkey | 1:800 | Jackson Immuno Research | 706-165-148 | AB_2340460 |
| Goat | Cy3 | Donkey | 1:800 | Jackson Immuno Research | 705-165-147 | AB_2307351 |
| Goat | Cy5 | Donkey | 1:800 | Jackson Immuno Research | 705-175-147 | AB_2340415 |
| Rabbit | Cy3 | Donkey | 1:800 | Jackson Immuno Research | 711-165-152 | AB_2307443 |
| Rabbit | Cy5 | Donkey | 1:800 | Jackson Immuno Research | 711-175-152 | AB_2340607 |
| Rat | Cy3 | Donkey | 1:800 | Jackson Immuno Research | 712-165-153 | AB_2340667 |
| Rat | Cy5 | Donkey | 1:800 | Jackson Immuno Research | 712-175-153 | AB_2340672 |

For wholemount preparations, each tissue piece was cut open along the mesenteric border and tautly pinned mucosa-side up on Sylgard 170 in ice-cold PBS. A paintbrush was used to gently remove any intestinal contents. Full-thickness tissue preparations were fixed in 4% PFA for 2 hours at 4 °C while shaking. After fixation and 3 washes in ice-cold PBS, tissue was pinned mucosa-side down on Sylgard 170, and fine forceps were used to peel the muscularis (smooth muscle layers and myenteric plexus (MP)) away from the underlying mucosa. The mucosa and muscularis layers were either used immediately or stored in PBS with 0.1% sodium azide at 4°C for up to 1 month. For immunostaining, tissue was cut into ∼2 cm long pieces and placed into WHO microtitration trays (International Scientific Supplies) with PBT. Tissue was then placed in primary antibodies (see Table 1) diluted in PBT overnight at 4°C while shaking. Tissue was washed 3 times in PBT, incubated in PBT containing secondary antibodies (see Table 2) for 2 hours at room temperature while shaking, washed twice in PBT, treated with PBS containing DAPI (1 mg/mL) for 5 minutes, and washed twice in PBS. For mounting, tissue was submerged in ddH_2_O, gently placed onto slides using paintbrushes under a dissection microscope, and allowed to briefly air dry before being coverslipped with Fluoromount-G (Southern Biotech).

### Image acquisition

Images were captured on a Leica SP8 confocal microscope using a 20x (NA 0.75) oil objective. Acquisition areas were selected, acquired, and stitched with LASX Navigator Mode (Leica). Images were captured as z-stacks, which were set at 3 μm intervals capturing the full depth of the area of interest, such as the MP. Images were subsequently processed with ImageJ/FIJI (NIH, Bethesda, MD) and Photoshop (Adobe).

### Image analysis

#### Depth projection

All image analyses were performed in ImageJ/FIJI (NIH, Bethesda, MD). Depth projection images of HuC/D labeling in the MP were generated and pseudocolored from z-stacks using the Temporal-Color Code macro in ImageJ/FIJI.

#### Conditional intensity function (CIF) analysis

Spatial probability maps were generated with CIF analysis as previously described^3^ with some modifications. Briefly, 800×800 μm maximum intensity projections of HuC/D+ neurons in MP wholemounts were processed with blur, threshold, and watershed functions. Analyze Particles was used to determine the XY coordinates of neurons in the tissue. Results were further analyzed with CIF algorithm, which assesses the number of neurons in a 2D grid around each neuron in the tissue. Results were smoothed with a Gaussian of 20 μm standard deviation. All analyses were performed in Python.

#### z-score difference analysis

Comparison of z-scores between neuronal data and synthetic data was performed as previously described^3^ with some modifications. Briefly, to determine the deviation of neuronal organization from complete spatial randomness over developmental time, XY coordinates of HuC/D+ neurons determined with Analyze Particles as above and assigned a z-score that reflects the deviation from the value expected with randomness. 500 samples of synthetic data were generated, and the mean z-score of these trials was assessed. In the generation of synthetic pseudorandom data, a minimum distance of 5 μm was imposed because the neuronal data did not overlap in space. To calculate the z-score difference, the z-score from 500 random samples was subtracted from the z-score of the data. All analyses were performed in Python.

#### Neuronal subtype counting

To count neuronal subtypes as an overall percentage of HuC/D+ neurons, image stacks of HuC/D immunolabeling in MP wholemount preparations were blurred and thresholded. Because HuC/D+ neurons in our tissue were tightly packed and not easily resolved, we approximated neuronal number from the overall area of HuC/D labeling. The HuC/D stack was maximally projected, and the Analyze Particles function was used to determine the total area labeled with HuC/D. To estimate the average area of a human fetal HuC/D+ enteric neuron, neuronal cell area was measured in multiple samples across different regions and ages, and approximately 138 μm^2^ was found to be the average neuronal area regardless of region or age (data not shown). Total HuC/D area was divided by this value to approximate the number of neurons in an image. To determine the percentage of HuC/D+ neurons that express a given subtype marker, stacks labeled with a subtype marker were blurred and combined with the processed HuC/D stack using the Image Calculator function. This result was maximally projected, and the number of cells expressing HuC/D and a given subtype marker, which were less densely packed and more easily resolved than HuC/D labeling, were counted with Analyze particles. This value was divided by the approximate number of HuC/D+ neurons to determine the overall representation of the subtype.

#### *Ex vivo* gastrointestinal (GI) motility assay and analysis

##### Ex vivo *GI motility monitor*

GI motility was assessed *ex vivo* using a setup adapted from platforms previously described.^15-17^ Tissue was placed in a dissection dish with Sylgard 170 and submerged in carbogenated Krebs solution as detailed above at 37 °C. The mesentery was cut away, and the tissue was moved to an organ bath above a heated water bath, which maintained the circulating carbogenated Krebs solution at 37 °C. The tissue was tautly pinned at the mesenteric border to the Sylgard 170 in the base of the chamber. After 10 minutes of acclimation, 10-minute videos were recorded at 3.75 frames/second using IC capture software (Imaging Source) and a high-resolution monochromatic firewire industrial camera (Imaging Source®, DMK41AF02) connected to a 2/3” 16 mm f/1.4 C-Mount Fixed Focal Lens (Fujinon HF16SA1) mounted above the organ bath. 30 minutes were captured at baseline followed by 30 minutes with the addition of 1 μM tetrodotoxin (TTX) (Alomone labs) to the circulating Krebs. A final 30 minutes of video were captured as the TTX solution was washed out. For the 20PCW sample, TTX was not used, and instead 60 minutes of video were captured at baseline. After completion of the experiment, tissue was collected, placed in ice-cold PBS, and processed for histology as described above. When the MP was removed, regions that did not contain a fully intact MP around their circumference were noted, and these regions were not subsequently analyzed for GI motility.

##### Spatiotemporal map (STM) analysis

STMs were generated from video recordings using VolumetryG9a software as previously described.^15^ For each region analyzed, two videos per condition (baseline, TTX, and washout) were assessed. STMs were visually inspected for motility patterns and evidence of distally propagating contractions. Speed and frequency of myogenic contractions, as indicated on the STM in Fig.2c, were manually measured with the line segment and measurement functions in ImageJ/FIJI for a 1 minute stretch of every STM containing myogenic ripples.

## Graphing and statistical analysis

All graphs and statistical analyses were generated with Prism 9 software (GraphPad). Each measurement represents a distinct human tissue sample. Given the low sample number for any given intestinal region and age, statistical outcomes were not reported in figures. Results of one-sided Tukey’s correction for multiple comparisons can be found in Supplementary Table 1.

## Data availability

Data generated and analyzed in this study are available upon reasonable request.

## Code availability

Code used to analyze these data are freely available at: https://github.com/druckmann-lab/EntericNervousSystemAnalysis. ^3^

## Acknowledgements

We thank Kaltschmidt lab members for experimental advice and discussions of results. In particular, we thank Julieta Gomez-Frittelli for feedback on the manuscript. We want to thank Shaul Druckmann and Aiden Wang for sharing analysis software before publication. We thank Subhamoy Das and Estelle Spear for setting up the *ex vivo* motility system in the Kaltschmidt lab. This work was supported by a Stanford Medical Scientist Training Program grant T32 GM007365-44 (L.B.D.), a Stanford Maternal and Child Health Research Institute Pilot Grant postdoctoral fellowship (L.L.), a Stanford Maternal and Child Health Research Institute Pilot Grant, the Wu Tsai Neurosciences Institute, the Stanford University Department of Neurosurgery, and a research grant from The Firmenich Foundation (J.A.K.).

## Author contributions

L.B.D. designed, performed, and analyzed all experiments and co-wrote the manuscript. L.L. collected tissue samples. A.M.P. organized tissue collection, contributed to discussions about experimental design and data analyses, and edited the manuscript. J.A.K. designed and supervised experiments and co-wrote the manuscript.

