## Supplemental information for "Anatomical and functional maturation of the mid-gestation human intestine"

### Extended data figure and movies

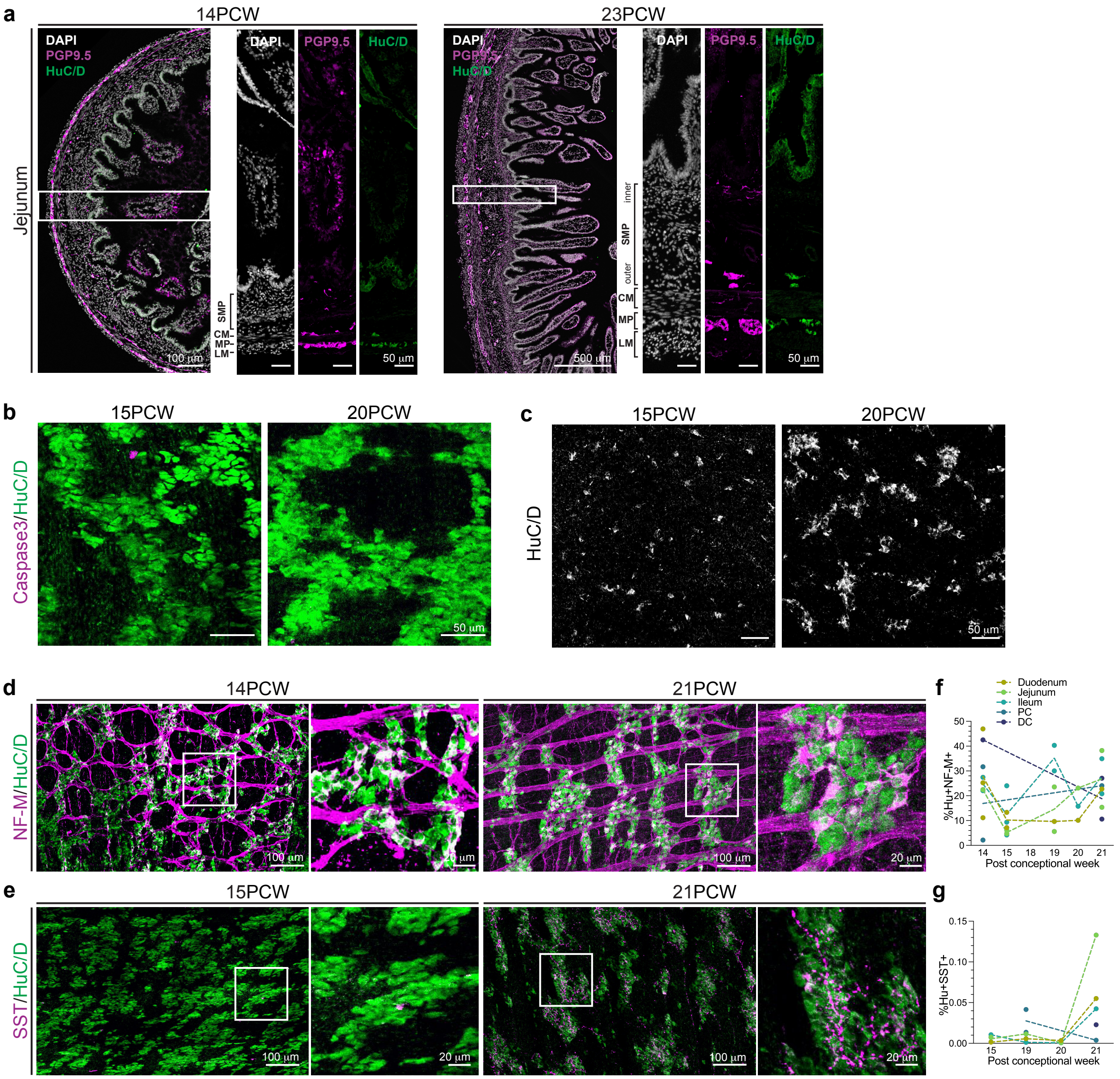


**Supplementary Figure 1 (Supporting Figure 1): Histological analysis of apoptosis, the submucosal plexus, and additional neuronal subtypes.** (**a**) Representative images of cross sections from the jejunum at 14 and 23 postconceptional weeks (PCW) with immunohistochemical (IHC) labeling against DAPI (white), pan-neuronal markers PGP9.5 (magenta) and HuC/D (green). White boxes indicate locations of higher magnitude insets. Brackets denote location of the submucosal plexus (SMP), circular muscle (CM), myenteric plexus (MP), and longitudinal muscle (LM). (**b**) Representative images of IHC labeling with apoptotic marker Caspase3 (magenta) and

HuC/D (green) in MP wholemount preparations at 15 and 20PCW. (**c**) Representative images of HuC/D labeling in wholemount preparation of the outer SMP in the small intestine at 15 and 20PCW. (**d,e**) IHC labeling in the jejunum MP with HuC/D (green) and sensory neuron marker NF-M (**d**, magenta) at 14 and 21PCW and interneuron marker SST (**e**, magenta) at 15 and 21PCW. White boxes indicate locations of higher magnitude insets. (**f,g**) Proportion of total HuC/D+ neurons positive for NF-M (**e**) or SST (**f**) (mean ± SEM) over developmental time. n = 1-3. Scale bars as indicated.

**Supplementary Movie 1 (Supporting Figure 2a,b): *ex vivo* motility of the 14PCW jejunum.** 10-minute video at 20x speed of a 30 mm segment of the 14 postconceptional week jejunum in the *ex vivo* gastrointestinal motility monitor. Tissue is oriented from proximal (left) to distal (right). Still images and accompanying spatiotemporal map for part of this video can be found in Figs.2a,b.

**Supplementary Movie 2 (Supporting Figure 2a): *ex vivo* motility of the 15PCW jejunum.** 10-minute video at 20x speed of a 30 mm segment of the 15 postconceptional week jejunum in the *ex vivo* gastrointestinal motility monitor. Tissue is oriented from proximal (left) to distal (right). Still image and accompanying spatiotemporal map for part of this video can be found in Fig.2a.

**Supplementary Movie 3 (Supporting Figure 2a): *ex vivo* motility of the 20PCW jejunum.** 10-minute video at 20x speed of a 30 mm segment of the 20 postconceptional week jejunum in the *ex vivo* gastrointestinal motility monitor. Tissue is oriented from proximal (left) to distal (right). Still image and accompanying spatiotemporal map for part of this video can be found in Fig.2a.

**Supplementary Movie 4 (Supporting Figure 2a,b): *ex vivo* motility of the 21PCW jejunum.** 10-minute video at 20x speed of a 30 mm segment of the 21 postconceptional week jejunum in the *ex vivo* gastrointestinal motility monitor. Tissue is oriented from proximal (left) to distal (right). Still images and accompanying spatiotemporal map for part of this video can be found in Figs.2a,b.

**Supplementary Movie 5 (Supporting Figure 2c): *ex vivo* motility of the 14PCW duodenum at baseline.** 10-minute video at 20x speed of a 25 mm segment of the 14 postconceptional week duodenum in the *ex vivo* gastrointestinal motility monitor prior to addition of tetrodotoxin. Tissue is oriented from proximal (left) to distal (right). Still image and accompanying spatiotemporal map for part of this video can be found in Fig.2c.

**Supplementary Movie 6 (Supporting Figure 2c):** ***ex vivo*** **motility of the 14PCW duodenum with tetrodotoxin.** 10-minute video at 20x speed of a 25 mm segment of the 14 postconceptional week duodenum in the *ex vivo* gastrointestinal motility monitor after the addition of 100 μM tetrodotoxin. Tissue is oriented from proximal (left) to distal (right). Still image and accompanying spatiotemporal map for part of this video can be found in Fig.2c.

**Supplementary Movie 7 (Supporting Figure 2c):** ***ex vivo*** **motility of the 21PCW duodenum at baseline.** 10-minute video at 20x speed of a 25 mm segment of the 21 postconceptional week duodenum in the *ex vivo* gastrointestinal motility monitor prior to addition of tetrodotoxin. Tissue is oriented from proximal (left) to distal (right). Still image and accompanying spatiotemporal map for part of this video can be found in Fig.2c.

**Supplementary Movie 8 (Supporting Figure 2c):** ***ex vivo*** **motility of the 21PCW duodenum with tetrodotoxin.** 10-minute video at 20x speed of a 25 mm segment of the 14 postconceptional week duodenum in the *ex vivo* gastrointestinal motility monitor after the addition of 100 μM tetrodotoxin. Tissue is oriented from proximal (left) to distal (right). Still image and accompanying spatiotemporal map for part of this video can be found in Fig.2c.
