## Supplementary Table 1 for "Anatomical and functional maturation of the mid-gestation human intestine"

**Fig.1f z-score difference**

| **Ordinary one-way ANOVA summary** | |
|---|---|
| F | 9.073 |
| P value | <0.0001 |
| P value summary | **** |
| Significant diff. among means (P < 0.05)? | Yes |
| R squared | 0.5645 |

| **ANOVA table** | **SS** | **DF** |
|---|---|---|
| Treatment (between columns) | 576.2 | 4 |
| Residual (within columns) | 444.6 | 28 |
| Total | 1021 | 32 |

| **Tukey's multiple comparisons test** | **Mean Diff.** | **95.00% CI of diff.** |
|---|---|---|
| Duodenum vs. Jejunum | 0.7727 | -5.433 to 6.978 |
| Duodenum vs. Ileum | -2.1 | -8.306 to 4.106 |
| Duodenum vs. PC | -9.585 | -16.04 to -3.126 |
| Duodenum vs. DC | -8.098 | -14.56 to -1.639 |
| Jejunum vs. Ileum | -2.873 | -9.078 to 3.333 |
| Jejunum vs. PC | -10.36 | -16.82 to -3.898 |
| Jejunum vs. DC | -8.87 | -15.33 to -2.411 |
| Ileum vs. PC | -7.485 | -13.94 to -1.026 |
| Ileum vs. DC | -5.998 | -12.46 to 0.4613 |
| PC vs. DC | 1.487 | -5.216 to 8.190 |

**Fig.1i %Hu+CalR+**

| **Ordinary one-way ANOVA summary** | |
|---|---|
| F | 6.289 |
| P value | 0.0005 |
| P value summary | *** |
| Significant diff. among means (P < 0.05)? | Yes |
| R squared | 0.6213 |

| **ANOVA table** | **SS** | **DF** |
|---|---|---|
| Treatment (between columns) | 237.9 | 6 |
| Residual (within columns) | 145 | 23 |
| Total | 382.8 | 29 |

| **Tukey's multiple comparisons test** | **Mean Diff.** | **95.00% CI of diff.** |
|---|---|---|
| 14 vs. 15 | -0.6355 | -6.545 to 5.274 |
| 14 vs. 18 | -1.561 | -6.990 to 3.868 |
| 14 vs. 19 | -3.309 | -8.427 to 1.809 |

| | | |
|---|---|---|
| 14 vs. 20 | -2.895 | -8.805 to 3.015 |
| 14 vs. 21 | -2.251 | -7.369 to 2.867 |
| 14 vs. 23 | -8.737 | -13.86 to -3.619 |
| 15 vs. 18 | -0.9255 | -7.106 to 5.255 |
| 15 vs. 19 | -2.673 | -8.583 to 3.237 |
| 15 vs. 20 | -2.259 | -8.867 to 4.348 |
| 15 vs. 21 | -1.615 | -7.525 to 4.294 |
| 15 vs. 23 | -8.102 | -14.01 to -2.192 |
| 18 vs. 19 | -1.748 | -7.176 to 3.681 |
| 18 vs. 20 | -1.334 | -7.514 to 4.847 |
| 18 vs. 21 | -0.69 | -6.119 to 4.739 |
| 18 vs. 23 | -7.176 | -12.60 to -1.748 |
| 19 vs. 20 | 0.4141 | -5.496 to 6.324 |
| 19 vs. 21 | 1.058 | -4.060 to 6.176 |
| 19 vs. 23 | -5.429 | -10.55 to -0.3104 |
| 20 vs. 21 | 0.6436 | -5.266 to 6.554 |
| 20 vs. 23 | -5.843 | -11.75 to 0.06721 |
| 21 vs. 23 | -6.486 | -11.60 to -1.368 |

**Fig.1j %Hu+nNOS+**

**Ordinary one-way ANOVA summary**

| | | |
|---|---|---|
| F | | 3.554 |
| P value | | 0.0116 |
| P value summary | * | |
| Significant diff. among means (P < 0.05)? | Yes | |
| R squared | | 0.4704 |

| **ANOVA table** | **SS** | **DF** |
|---|---|---|
| Treatment (between columns) | 1590 | 6 |
| Residual (within columns) | 1790 | 24 |
| Total | 3379 | 30 |

| **Tukey's multiple comparisons test** | **Mean Diff.** | **95.00% CI of diff.** |
|---|---|---|
| 14 vs. 15 | -12.26 | -32.51 to 7.992 |
| 14 vs. 18 | -5.764 | -23.30 to 11.77 |
| 14 vs. 19 | 9.962 | -7.576 to 27.50 |
| 14 vs. 20 | 9.372 | -10.88 to 29.62 |
| 14 vs. 21 | -4.645 | -22.18 to 12.89 |
| 14 vs. 23 | 4.699 | -12.84 to 22.24 |
| 15 vs. 18 | 6.495 | -13.76 to 26.75 |
| 15 vs. 19 | 22.22 | 1.970 to 42.47 |
| 15 vs. 20 | 21.63 | -1.009 to 44.27 |

| | | |
|---|---|---|
| 15 vs. 21 | 7.614 | -12.64 to 27.86 |
| 15 vs. 23 | 16.96 | -3.292 to 37.21 |
| 18 vs. 19 | 15.73 | -1.812 to 33.26 |
| 18 vs. 20 | 15.14 | -5.114 to 35.39 |
| 18 vs. 21 | 1.119 | -16.42 to 18.66 |
| 18 vs. 23 | 10.46 | -7.074 to 28.00 |
| 19 vs. 20 | -0.5893 | -20.84 to 19.66 |
| 19 vs. 21 | -14.61 | -32.14 to 2.931 |
| 19 vs. 23 | -5.262 | -22.80 to 12.28 |
| 20 vs. 21 | -14.02 | -34.27 to 6.233 |
| 20 vs. 23 | -4.673 | -24.92 to 15.58 |
| 21 vs. 23 | 9.345 | -8.193 to 26.88 |

**Fig.2d Frequency**

| **Two-tailed paired t test summary** | | **P value** | **Mean of Baseline** |
|---|---|---|---|
| 14 | | | |
| 15 | No | | 0.67859 |
| 20 | | | |

**Fig.2e Speed**

| **Two-tailed paired t test summary** | | **P value** | **Mean of Baseline** |
|---|---|---|---|
| 14 | | | 0.2748 |
| 15 | | 0.197379 | 0.347 |
| 20 | | | 0.1702 |

**Supplementary Fig.1f %Hu+NFM+**

| **Ordinary one-way ANOVA summary** | | |
|---|---|---|
| F | | 2.71 |
| P value | | 0.0732 |
| P value summary | ns | |
| Significant diff. among means (P < 0.05)? | No | |
| R squared | | 0.4364 |

| **ANOVA table** | **SS** | **DF** |
|---|---|---|
| Treatment (between columns) | 696.3 | 4 |
| Residual (within columns) | 899.3 | 14 |
| Total | 1596 | 18 |

| **Tukey's multiple comparisons test** | **Mean Diff.** | **95.00% CI of diff.** |
|---|---|---|
| 14 vs. 15 | 17.58 | -0.6627 to 35.81 |

| | | |
|---|---|---|
| 14 vs. 19 | 7.082 | -11.16 to 25.32 |
| 14 vs. 20 | 10.56 | -7.675 to 28.80 |
| 14 vs. 21 | 2.832 | -12.96 to 18.63 |
| 15 vs. 19 | -10.49 | -30.88 to 9.897 |
| 15 vs. 20 | -7.012 | -27.40 to 13.38 |
| 15 vs. 21 | -14.74 | -32.98 to 3.495 |
| 19 vs. 20 | 3.481 | -16.91 to 23.87 |
| 19 vs. 21 | -4.249 | -22.49 to 13.99 |
| 20 vs. 21 | -7.731 | -25.97 to 10.51 |

**Supplementary Fig.1g %Hu+SST+**

| **Ordinary one-way ANOVA summary** | | |
|---|---|---|
| F | 2.429 | |
| P value | 0.1204 | |
| P value summary | ns | |
| Significant diff. among means (P < 0.05)? | No | |
| R squared | 0.3985 | |
| **ANOVA table** | **SS** | **DF** |
| Treatment (between columns) | 0.006795 | 3 |
| Residual (within columns) | 0.01026 | 11 |
| Total | 0.01705 | 14 |
| **Tukey's multiple comparisons test** | **Mean Diff.** | **95.00% CI of diff.** |
| 15 vs. 19 | -0.005592 | -0.07578 to 0.0646[illegible] |
| 15 vs. 20 | 0.004311 | -0.07073 to 0.0793[illegible] |
| 15 vs. 21 | -0.04552 | -0.1126 to 0.02160 |
| 19 vs. 20 | 0.009903 | -0.06029 to 0.0800[illegible] |
| 19 vs. 21 | -0.03993 | -0.1016 to 0.02172 |
| 20 vs. 21 | -0.04983 | -0.1169 to 0.01729 |

| MS | F (DFn, DFd) | P value |
|---|---|---|
| 144.1 | F (4, 28) = 9.073 | P<0.0001 |
| 15.88 | | |

| Summary | Adjusted P Value |
|---|---|
| ns | 0.9961 |
| ns | 0.8593 |
| ** | 0.0015 |
| ** | 0.0086 |
| ns | 0.664 |
| *** | 0.0006 |
| ** | 0.0035 |
| * | 0.017 |
| ns | 0.0785 |
| ns | 0.9659 |

| MS | F (DFn, DFd) | P value |
|---|---|---|
| 39.64 | F (6, 23) = 6.289 | P=0.0005 |
| 6.304 | | |

| Summary | Adjusted P Value |
|---|---|
| ns | 0.9998 |
| ns | 0.9642 |
| ns | 0.394 |

| | |
|---|---|
| ns | 0.6963 |
| ns | 0.7871 |
| *** | 0.0002 |
| ns | 0.9989 |
| ns | 0.7654 |
| ns | 0.9211 |
| ns | 0.9719 |
| ** | 0.0032 |
| ns | 0.9395 |
| ns | 0.9916 |
| ns | 0.9996 |
| ** | 0.0047 |
| ns | >0.9999 |
| ns | 0.9933 |
| * | 0.0327 |
| ns | 0.9998 |
| ns | 0.054 |
| ** | 0.0071 |

| **MS** | **F (DFn, DFd)** | **P value** |
|---|---|---|
| 265 | F (6, 24) = 3.554 | P=0.0116 |
| 74.57 | | |

| **Summary** | **Adjusted P Value** |
|---|---|
| ns | 0.4729 |
| ns | 0.935 |
| ns | 0.5457 |
| ns | 0.7499 |
| ns | 0.9765 |
| ns | 0.9751 |
| ns | 0.9417 |
| * | 0.025 |
| ns | 0.0678 |

| | |
|---|---|
| ns | 0.8845 |
| ns | 0.1442 |
| ns | 0.0997 |
| ns | 0.2413 |
| ns | >0.9999 |
| ns | 0.4897 |
| ns | >0.9999 |
| ns | 0.1482 |
| ns | 0.9571 |
| ns | 0.3206 |
| ns | 0.9883 |
| ns | 0.6156 |

| Mean of +Ttx | Difference | SE of difference |
|---|---|---|
| 0.1202 | 0.1227 | -0.002503 |
| 0.1373 | 0.1298 | 0.007481 |

| Mean of +Ttx | Difference | SE of difference |
|---|---|---|
| 0.2715 | 0.003246 | |
| 0.4338 | -0.08678 | 0.04561 |

| MS | F (DFn, DFd) | P value |
|---|---|---|
| 174.1 | F (4, 14) = 2.710 | P=0.0732 |
| 64.23 | | |

| Summary | Adjusted P Value |
|---|---|
| ns | 0.0613 |

| | |
|---|---|
| ns | 0.746 |
| ns | 0.4087 |
| ns | 0.979 |
| ns | 0.5187 |
| ns | 0.8178 |
| ns | 0.1418 |
| ns | 0.9824 |
| ns | 0.9468 |
| ns | 0.6836 |

| MS | F (DFn, DFd) | P value |
|---|---|---|
| 0.002265 | F (3, 11) = 2.429 | P=0.1204 |
| 0.0009325 | | |

| Summary | Adjusted P Value |
|---|---|
| ns | 0.9949 |
| ns | 0.998 |
| ns | 0.2314 |
| ns | 0.9731 |
| ns | 0.2638 |
| ns | 0.1737 |
